## Supplementary figures and images for "Effects of short-term isolation on vocal and non-vocal social behaviors in prairie voles"

### Supplemental Figures

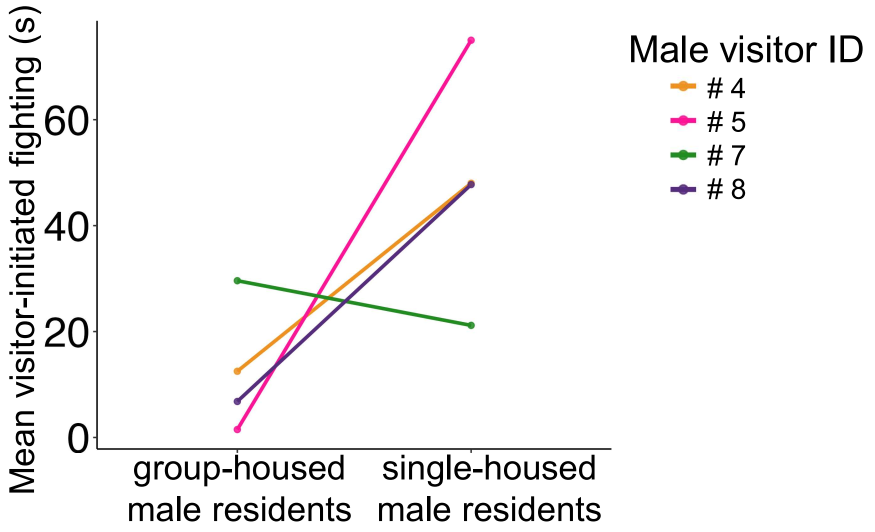

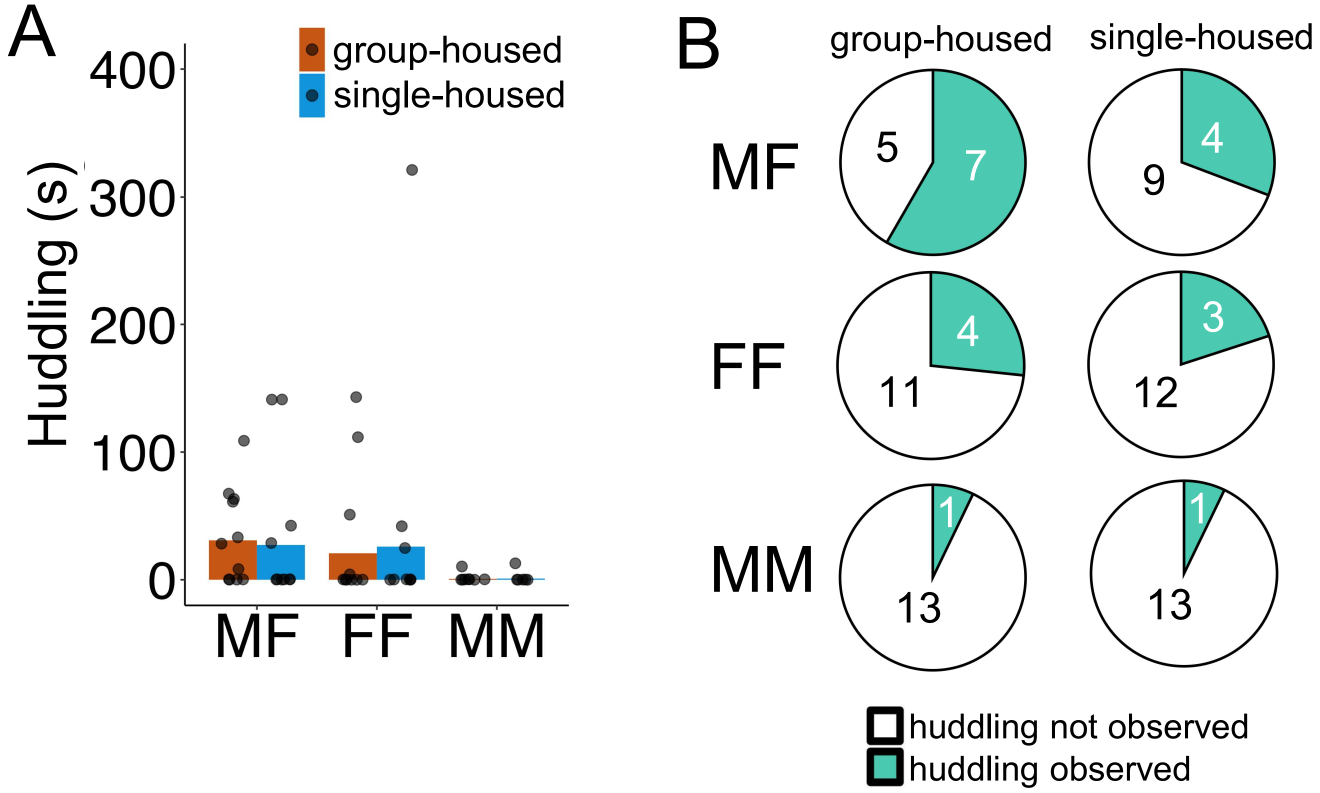
